# Patterns of genomic variation in the opportunistic pathogen *Candida glabrata* suggest the existence of mating and a secondary association to the human host

**DOI:** 10.1101/105791

**Authors:** Laia Carreté, Ewa Ksiezopolska, Cinta Pegueroles, Emilia Gómez-Molero, Ester Saus, Susana Iraola-Guzmán, Damian Loska, Oliver Bader, Cecile Fairhead, Toni Gabaldón

## Abstract

*Candida glabrata* is an opportunistic fungal pathogen that ranks as the second most common cause of systemic candidiasis. Despite its genus name, this yeast is more closely related to the model yeast *Saccharomyces cerevisiae* than to other *Candida* pathogens, and hence its ability to infect humans is thought to have emerged independently. Morover, *C. glabrata* has all the necessary genes to undergo a sexual cycle, but it is considered an asexual organism due to the lack of direct evidence of sexual reproduction. Here, we assessed genomic and phenotypic variation across 33 globally-distributed *C. glabrata* isolates. We cataloged extensive copy number variation, which particularly affects genes encoding cell-wall associated proteins, including adhesins. The observed level of genetic variation in *C. glabrata* is significantly larger than that found in *Candida albicans*. This variation is structured in seven deeply divergent clades, which show recent geographical dispersion and large within-clade genomic and phenotypic differences. We show compelling evidence of recent admixture between differentiated lineages, and of purifying selection on mating genes, which provide fist evidence for the existence of a sexual cycle in this yeast. Altogether, our data point to a recent global spread of previously genetically isolated populations and suggest that humans are only a secondary niche for this yeast.

## Introduction

The prevalence of infections by opportunistic pathogens (i.e. candidiasis) is increasing, partly due to recent medical progress enabling the survival of susceptible individuals (Brown et al. 2012). Main prevalent agents of candidiasis comprise three *Candida* species: *Candida albicans, Candida glabrata* and *Candida parapsilosis*, generally in this order (Diekema et al. 2012). Phylogenetically, these species are only distantly related. *C. glabrata* belongs to the *Nakaseomyces* clade, a group which is more closely related to the baker's yeast *Saccharomyces cerevisiae* than to *C. albicans* or *C. parapsilosis* (Gabaldón et al. 2013). Furthermore, both *C. glabrata* and *C. albicans* have closely related non-pathogenic relatives, hence the ability to infect humans in these two lineages must have originated independently (Gabaldón and Carreté 2016; Gabaldón et al. 2016). Genome sequencing of non-pathogenic and mildly-pathogenic relatives of *C. glabrata* has enabled tracing the genomic changes that correlate with the evolutionary emergence of pathogenesis in the *Nakaseomyces* group (Gabaldón et al. 2013). These analyses revealed that the ability to infect humans has likely emerged at least twice independently in the *Nakaseomyces*, coinciding with parallel expansions of the encoded repertoire of cell-wall adhesins. Thus, increased - or more versatile - adherence may be implicated in the evolutionary emergence of virulence potential towards humans. In contrast, other virulence-related characteristics had a more ancient origin within the clade and were also found in environmental relatives. Our understanding of the evolution of *C. glabrata* at the species level is limited to analyses of natural variation restricted to a few loci (Dodgson et al. 2005, 2003; Brisse et al. 2009). These studies have shown the existence of genetically distinct clades and generally suggested clonal, geographically structured populations. Geographically structured populations are also found in *C. albicans*, which is tightly associated to humans and which can undergo a parasexual cycle (Fundyga et al. 2002; Hirakawa et al. 2015; Tavanti et al. 2005), and *S. cerevisiae*, which has been domesticated and which can undergo a full sexual cycle, usually involving self-mating (Cromie et al. 2013; Liti et al. 2009). *C. glabrata* has been described as an asexual species despite the presence of homologs of *S. cerevisiae* genes involved in mating (Fabre et al. 2005). In order to shed light on the recent evolution of this important opportunistic pathogen we analyzed the genomes and phenotypes of 33 different clinical and colonizing *C. glabrata* isolates sampled from different human body sites and globally distributed locations, and chosen within genotyped.collections so as to be representative of previously explored population structure (Enache-Angoulvant et al. 2010; Brisse et al. 2009) (Supplementary Table 1). This sampling includes the extensively studied BG2 strain, as well as three pairs of strains, each isolated from a single patient.

## Results and discussion

### High levels of genetic diversity and lack of strong geographical structure

Using a read-mapping strategy against the available reference genome (Dujon et al. 2004a) we cataloged single nucleotide polymorphisms (SNPs) and copy number variations (CNVs) (see Materials and Methods). Overall we detected a range of 4.66-6.56 SNPs/Kb per strain when compared against the reference, a range of 0.04-7.23 SNPs/Kb between pairs of strains from different patients, and a range of 0.05– 0.07 SNPs/Kb between strains from the same patient. This indicates that patients are colonized by a single *C. glabrata* population that can be found at different body sites. We used Multiple Correspondence Analysis (MCA), Maximum Likelihood (ML) phylogenetic reconstruction, and model-based clustering, to establish the main relationships between all sequenced strains (Figure 1). Overall, these analyses support the existence of seven major clades, hereafter referred as clade I through clade VII. Of note, model-based clustering of genetic variation provided strong indication of recent genetic admixture between different clades. In particular individuals of clade II may have undergone extensive recombination with clade I, as suggested by both model-based clustering (Figure 1c) and pairwise differences in SNP density (Supplementary Figure S1). Phylogenetic reconstruction and fixation indices (FST) indicate that most clades diverged deeply within the *C. glabrata* lineage. Genetic distance between the two most distant clades (clade I and clade VII, 6.59-7.22 SNPs/Kb) is only slightly higher than that between the most closely related ones (clade I, clade II and clade III, 4.48-6 SNPs/Kb, Supplementary Figure S2), but up to two orders of magnitude larger than the amount of genetic divergence within clades (0.03-0.29 SNPs/Kb for all clades except clade V, with 4.37-4.68 SNPs/Kb). Comparatively, the level of variation within some of the *C. glabrata* clades is similar to the amount of genetic variation among distant clades in human associated *C. albicans* (average of 3.7 SNPs/Kb) (Hirakawa et al. 2015). Thus the genetic diversity of human-associated *C. glabrata* strains is significantly larger than that found in *C. albicans*. Most clades were present across distant locations and in different body sites, but they were generally enriched in one of the two mating types (Figure 1, Supplementary Figure S3). After these analyses were performed, the genome from the pyruvate-producing strain *C. glabrata* CCTCC M202019 was published (Xu et al. 2016). This strain is said to have been isolated from fertile soil, and thus it represents the only available genome sequence from a strain not isolated form the human body. Our re-analysis of this strain (see Materials and Methods) identifies relatively few SNPs when compared against the reference CBS138 (1026 SNPs, 0.08 SNPs/Kb), which situates this strain within clade V, and as the most similar to the reference genome. Current lack of a strong geographical structure of deeply divergent clades suggests recent global migration of human-associated *C. glabrata* strains. Finally, the appearance of a strain isolated from soil within a clade of human-associated strains suggests strains with similar genetic backgrounds can colonize humans and the environment.

**Figure 1.**
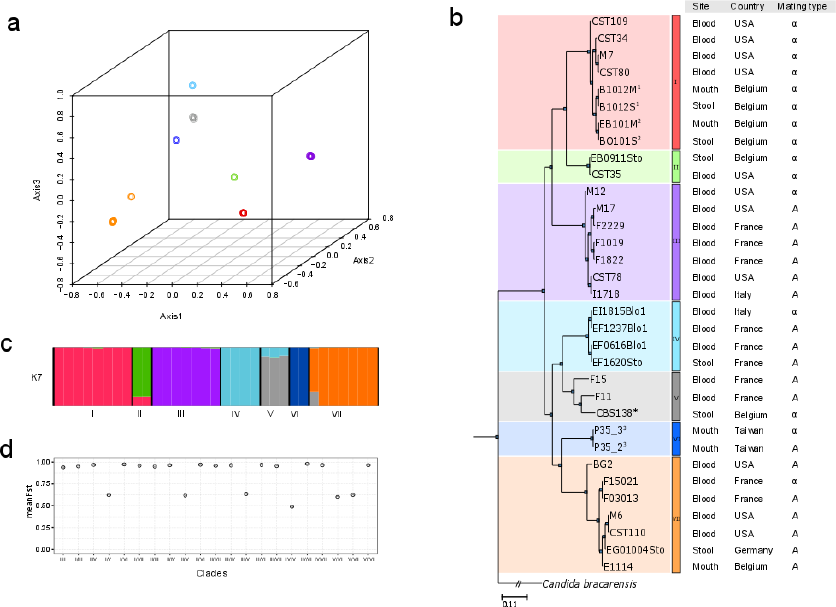
Distribution and population structure of the 33 strains of *C. glabrata* based on SNP data analysis. a) 3Dscatterplot of the multiple correspondence analysis (MCA), in which the different colors designate the seven clades detected. b) Phylogenetic tree computed using a maximum-likelihood approach. Super-indices indicate pairs of strains in which the two originate from the same patient (different body site, or different isolation date, see Supplementary Table S1 for more details). Clades from I to VII were designated using the same colors as in A. c) Population admixture using STRUCTURE software with K=7 using the same colors as A. d) Mean F_ST_ for all pairwise comparisons between the seven clades. Fisher test was used to analyze association with geographical structure (*P*(country-clade)=0.006), body site of isolation (*P*(site-clade)=0.157) and mating type (*P*(mating-clade) =6.064e-05).

### Evidence for genomic recombination between distinct clades

Using depth-of-coverage analyses, we detected a total of 46 deleted, and 62 duplicated genes (Figure 2, Supplementary Table S2). Of these we experimentally confirmed a deletion covering three genes (see Supplementary Figure S4). A significant fraction of the deleted (45.65%) or duplicated (41.94%) genes encoded GPI-anchored adhesin-like proteins, as compared to the 1.3% that this functional category represents over the entire genome (de Groot et al. 2013). Taken together, analysis of biallelic SNPs, flow cytometry, and electrophoretic karyotyping indicate that all analyzed strains are haploid, albeit with variations in total DNA content, and chromosome numbers and lengths (Supplementary Figures S5-7). Depth of coverage analysis revealed aneuploidies involving a whole duplication of chromosome E, whose presence is interspersed in different clades, and one strain carrying a partial aneuploidy of chromosome G (Figure 2, Supplementary Figure S8).

**Figure 2.**
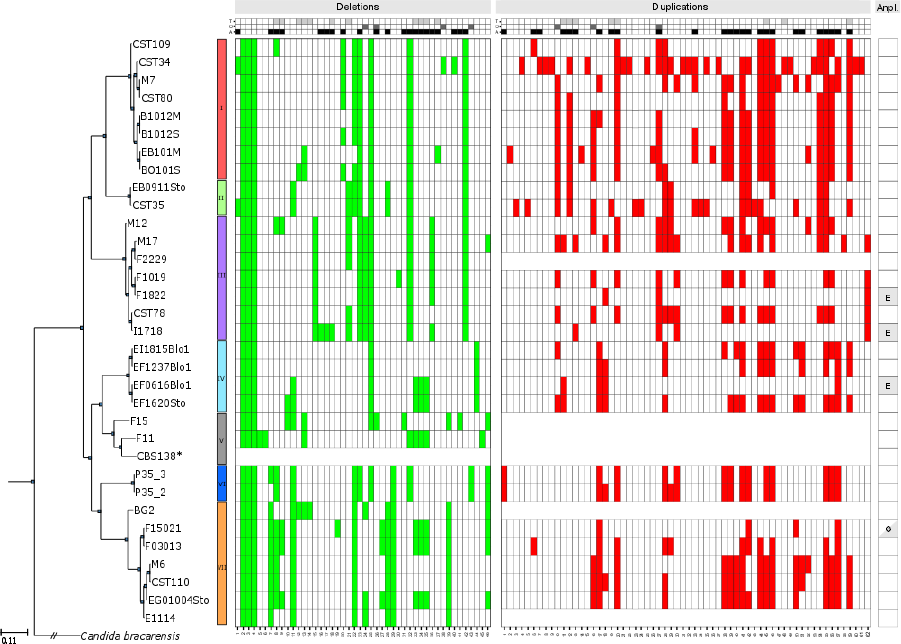
Heatmap showing the deletions, duplications, and aneuploidies (Anpl.) detected in the analyzed strains of *C.glabrata* sorted by clade. Reference (CBS138) and chromosomes with aneuploidies in affected strains (see below) or genomes with unstable coverage are not shown. Heatmap on the top of the figure designates gene information; light grey, grey and black represent genes in a tandem duplication (T), orphan genes (O) and genes encoding GPI-anchored adhesin-like proteins (A) respectively. The heatmap colored in green designates the 46 genes affected by deletions and the heatmap colored in red designate the 62 genes affected by duplications. Aneuploidies are indicated with a light grey background with the letter of chromosome affected. Fisher test was used to test the significant enrichment in genes encoded GPI-anchored adhesin-like proteins (p-value < 1.44e-26 and 5.16e-31 in deletions and duplications respectively), orphan proteins (p-value < 0.051 and 0.436 in deletions and duplications respectively) and genes in a tandem duplication (p-value < 1.474e-09 and 3.02e-05 in deletions and duplications respectively). See Supplementary Table S2 for the complete list of genes affected.

Aneuploid chromosomes had similar numbers of predicted heterozygous SNPs as other chromosomes when a diploid model was enforced in the SNP calling process (Supplementary Figure S9, see Materials and Methods), suggesting the extra chromosomes diverged recently. Although all aneuploidies affect genes related to drug resistance, the aneuploid strains had normal sensitivity to tested antifungals (see below). Interestingly, our sequencing data indicated that amajor duplication of chromosome J occurred spontaneously while growing one strain (F2229) in rich medium and in the absence of antifungals, as it was present only in about 50% of the cells at the time of sequencing (see Supplementary Figure S10). This underscores the plasticity of *C.glabrata* genome even under laboratory conditions (Bader et al. 2012). Using a *de novo* assembly strategy we found 20 different large re-arrangements grouped in 17 conformations and affecting 26 different strains, including fourteen translocations and three inversions (Figure 3), some of which confirmed previous reports based on electrophoretic karyotyping and comparative genome hybridization (Muller et al. 2009). The distribution of most CNVs and large-scale re-arrangement agreed with the above defined clades. However, we found several striking cases of deletions, and large re-arrangements which were shared between distantly related isolates (Figure 3).

**Figure 3.**
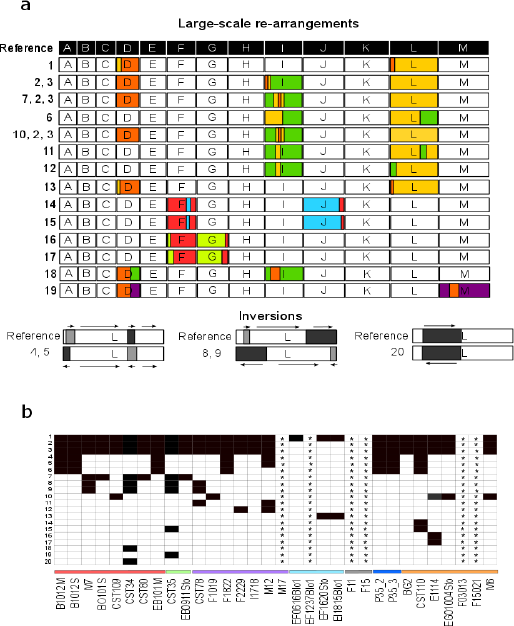
Chromosomal re-arrangements. a) Diagram showing the 20 different large re-arrangements found in*C.glabrata.* Re-arrangements are grouped in seventeen different conformations, including fourteen translocations and three inversions. Chromosomes are indicated in letters from A to M and independent rearrangements are designated by different colors. Arrows near the inversion plots indicate relative orientation of the indicated fragments. b) Heatmap showing the distribution of annotated rearrangements (1-20) and the 26 affected strains. Asterisks “*” indicate strains not included in the analysis due to high fragmentation of the assembly.

Given the close correspondence of the predicted boundaries of some re-arrangements we considered unlikely that these patterns emerged independently, and suspected the existence of recombination events between distinct lineages, which would further support the existence of admixture reported above. Detailed analyses of nineteen deletions shared by distant strains strongly suggest that seventeen of them are the result of genetic exchange mediated through genomic recombination (Figure 4, see Supplementary Figure S11).

**Figure 4.**
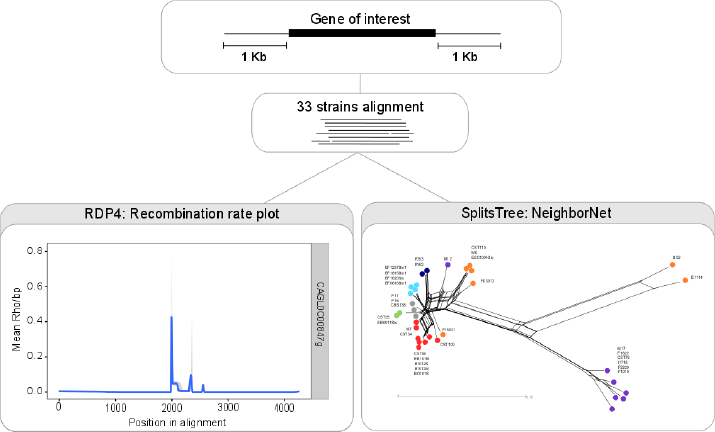
Recombination analyses. Analysis of the region surrounding the deletion in *CAGL0C00847g* gene is shown as an example. First, we selected regions containing the gene of interest and 1Kb flanking regions in the 33 strains. Second, we estimated recombination rates (rho/bp) in the selected regions and phylogenetic networks (see Materials and Methods). Bottom left) plot showing recombination rates along the genomic region. Values higher than 0.05 indicate a recombination hotspot. Bottom right) NeighborNet splits network showing gene flow between strains and the phylogenetic signal in the region. Clades are indicated as dots of different colors. Strains of different clades cluster together, suggesting a much closer genetic relationship than expected from the genome-wide analysis, which is indicative of recombination between different clades.

### Genes involved in mating are evolutionary constrained at the species level

The above results suggest that mating does occur in *C. glabrata*. If mating has played a role in *C.glabrata* adaptation, we expect these genes to show hallmarks of selective constraints at the species level. We assessed levels of genetic variation in *C. glabrata* genes, and compared these with those obtained here (see Materials and Methods, Supplementary Table S3) from re-analyzing published data in *C. albicans* and *S. cerevisiae*, which show parasexual and sexual cycles, respectively. At the genome-wide level the three species show overall similar levels of constraints (Figure 5).

**Figure 5:**
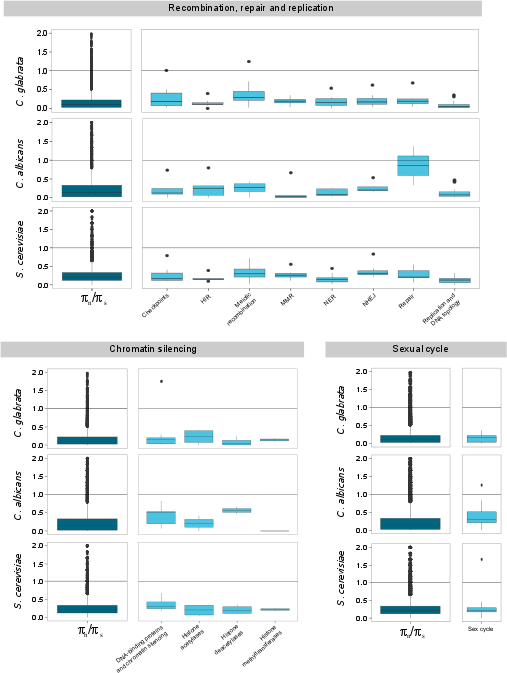
Ratio of non-synonymous and synonymous nucleotide diversity (πN/πS) in genes involved in mating and recombination in *C. glabrata,* and in their one-to-one orthologs in *C. albicans* and *S. cerevisiae*. Dark blue plots show overall πN/πS values in each category and light blue plots show specific groups of genes included in each category. The most distant outliers are not shown, as the length of the Y axis was limited to 2.

We next focused on three different classes of genes involved in mating and recombination: i) genes involved in the firsts steps of the sexual cycle (Fabre et al. 2005) ii) genes involved in chromatin silencing of sexual genes and regulation of mating-type cassettes (Fabre et al. 2005) and iii) genes involved in replication, repair and recombination (Richard et al. 2005) (Figure 5). All three classes showed signatures of constrains in the three species. Of note, while some classes are more constrained than others, similar patterns are observed in all species. For instance, genes involved in meiotic recombination and repair had signs of relaxation of selection as compared to genes involved in other cellular processes, whereas genes involved in mating or other processes of the sexual cycle are more constrained. We searched for genes showing an excess of non-synonymous variations in *C. glabrata* as compared to *S. cerevisiae* and *C. albicans* (Supplementary Table S4). This uncovered the orthologs of *S. cerevisiae* genes *ECS1, MEI4, REC14*, and *RAD9*, involved in silencing, meiotic double-strand break formation, meiotic recombination, and DNA damage repair, respectively. Importantly, *Ecs1p* interacts with Sir4p, andis involved in telomere silencing of the *HML* and *HMR* cassettes (Andrulis et al. 2002) in *S. cerevisiae*. Altogether, our results show that *C. glabrata* genes involved in mating and meiosis have comparable levels of selective constraints as those found in *C.albicans* and *S. cerevisiae*, providing support for the existence of a sexual or parasexual cycle in *C. glabrata*. Of note, the anomalous excess of non-synonymous mutation in the *C. glabrata* ortholog *ESC1*, suggestive of a recent functional shift, may provide a clue for the observed differences in silencing of mating loci between *S. cerevisiae* and *C.* glabrata (Muller et al. 2008).

### Illegitimate mating-type switching

Consistent with earlier observations (Brisse et al. 2009), our data supports the existence of mating-type switching, albeit very limited, in *C. glabrata* populations. Similar to *S. cerevisiae*, the *C.glabrata* genome encodes the two mating types (**a** and alpha) in three different loci called *MTL1 (MAT)*, *MTL2 (HMR)* and *MTL3 (HML)*, and the *HO* gene which encodes the endonuclease responsible for gene conversion based mating-type switching. *MTL2* and *MTL3* encode **a** and alpha information, respectively, and they are close to telomeres. The *MTL1* locus encodes either **a** or alpha and this information determines the mating type identity of the cell. Analysis of mating type loci revealed eight strains showing gene conversion events of four different types (Figure 6, Supplementary Figure S12). In three cases, a normal conversion event at *MTL1* switched the mating type from **a** to alpha. The five remaining cases represent cases of aberrant conversions. In one case, **a** to alpha switching at *MTL1* is accompanied by illegitimate conversion at *MTL2* resulting in a triple alpha strain. In three cases, illegitimate *MTL2* conversion occurred in the apparent absence of *MTL1* switching. A final case represents illegitimate conversion of *MTL3* in the apparent absence of *MTL1* switching leading to a triple **a** strain. In all aberrant cases, the correspondence of the conversion track with an *HO* cutting site strongly suggest that this switching is mediated by illegitimate cuts in *MTL2* or *MTL3*. These results show that aberrant conversions previously induced experimentally (Boisnard et al. 2015) occur in natural populations.

**Figure 6.**
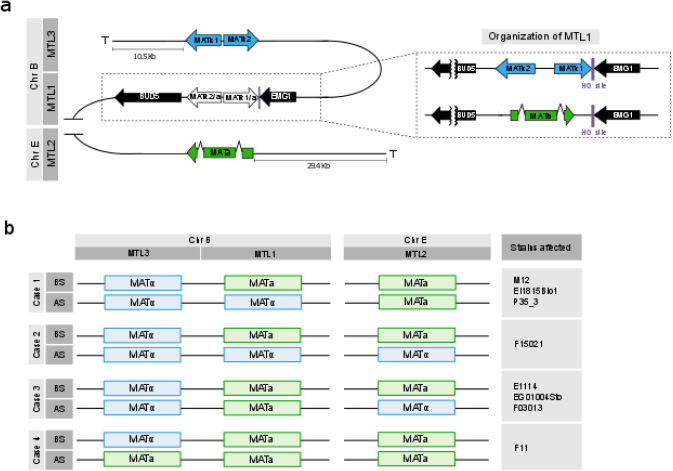
Mating type loci of *C. glabrata*. a) Organization of mating type loci in *C. glabrata*:*MTL1* in white, *MTL2* in green and *MTL3* in blue. *MTL1* is shown enlarged on the right, encoding either **a** or alpha type genes. b) Diagram of the four cases of mating type switching events likely to have occurred in sequenced strains. BS: Before switching. AS: After switching.

### Genomic plasticity enables large phenotypic differences between and within clades

To assess whether the observed genomic plasticity was reflected at the phenotypic level, we measured several relevant phenotypes of the sequenced strains. Specifically, we tested biofilm formation properties, antifungal drug susceptibility, and we measured growth under stress conditions such as high or low pH, high temperature, and presence of DTT, sodium chloride, or hydrogen peroxide (Materials and Methods, Supplementary Figure S13, and Supplementary table S5). Most conditions showed important differences between strains of the same clades. In fact, for most conditions and most clades, intra-clade variation was of a similar range as inter-clade variation. We surveyed private mutations, that is SNPs and CNVs present in a single strain of a clade (Supplementary Table S6). This resource may help identifying the genetic bases of phenotypic differences in strains behaving drastically different from their close relatives, as well as identifying common mutations in distant strains that show similar behaviors in a given condition. Three strains (M6, M7, M17) showed reduced sensitivity to one or more drugs, among eight tested (Supplementary Table S5). Each carried a unique, private mutation in *PDR1* (leading to amino acid exchanges I390K in M6, I378T in M7, and N306S in M17), a known regulator of pleiotropic drug response (Tsai et al. 2006). Recently it has been claimed that prevalent mutations in the mismatch repair gene *MSH2* found in clinical isolates promote drug resistance through a mutator phenotype (Healey et al. 2016). Fifteen (45%) of the 33 analyzed strains carry non-synonymous SNPs in that gene, which include only one (M17) of the three strains with reduced antifungal sensitivity. These 15 strains carried four different *MSH2* variants, of which two correspond to variants previously proposed to be loss-of-function mutator genotypes (V239L/A942T and V239L). However, these SNPs were shared by all strains in the same clade, correspond to fixed mutations in other yeast species, and strains with these genotypes did not show unusual patterns of non-synonymous or synonymous variations (see Supplementary Table S7). Altogether our results suggest that these mutations represent natural genetic variation and are likely not related to a mutator phenotype. We next focused on differences on adherence properties, a virulent trait that may vary depending on the repertoire of proteins attached to the cell-wall. We noted that three strains showed high (F03013) or moderately high (CST35, F15021) ability to form biofilms on polystyrol (Supplementary Figure S20). These three biofilm forming strains shared independent duplications of *PWP4* and deletions of *AWP13*, two GPI-anchored adhesins (de Groot et al. 2013). Although these and other genotype-phenotype relationships enabled by the current dataset can provide useful hints, further experiments are needed to assess what genomic alterations underlie a given phenotypic variation.

## Conclusions

Our results show that human-associated *C. glabrata* isolates belong to (at least) seven genetically distinct clades, some of which present levels of genetic diversity comparable to those found in the global *C. albicans* population. In addition, the absence of a strong geographical structure and the deep genetic divergence between the clades suggests a model of ancient geographical differentiation with recent global dispersion, likely mediated by humans. This recent dispersion has likely put in contact, for the first time, *C. glabrata* clades that had been separated over a long period of time. Importantly, our results show that this recent admixture has resulted in genetic exchange between distinct clades. The existence of some form of sexual cycle is also strongly supported by similar patterns of evolutionary constraints in reproduction related genes in *C. glabrata, C. albicans,* and *S.cerevisiae*. Our results are consistent with previous reports of successful mating-type switching in *MTL1* from **a** to alpha, but also reveal frequent illegitimate recombination at the other *MTL* loci. Importantly, the illegitimate recombinations most likely result from cutting of the *HO* endonuclease sites present in *MTL2* and *MTL3*, which are generally not targeted in *S. cerevisiae*. This difference may relate to the conformational or epigenetic status of these genomic regions. Our finding of an excess of non-synonymous variation of the *C. glabrata* ortholog of *ESC1*, encoding a protein that participates in telomere silencing, may provide the first clue to this fundamental difference, as a functional shift in this protein may have directly impacted the structural or epigenetic organization of *C. glabrata* telomeres and subtelomeric regions.

We report extensive phenotypic and genetic variation, even between closely related strains, indicating fast evolutionary dynamics and a potential for fast adaptation in *C. glabrata*. Genetic variation particularly affects cell-wall proteins involved in adhesion. A species-specific increase in adhesins has been purported as a key step in the emergence of the ability to infect humans in the *C.glabrata* lineage (Gabaldón et al. 2013). Our finding of a highly dynamic genetic repertoire of adhesins and large differences in adhesion capabilities, suggests that there is a large degree of standing variation of this trait, which may be the subject of ongoing directional selection. Most of these genes are encoded in subtelomeric regions. The above mentioned difference in *ESC1* could also be related to such dynamism. Indeed, null mutants of *S. cerevisiae ESC1* show higher chromosome instability and increased transposable elements transposition. Thus, it is tempting to speculate that a functional shift in *ESC1* may have impacted both mating-type switching and the dynamics of subtelomeric genes, and could also be related to the plasticity of chromosomal structure in *C. glabrata*.

## Materials and Methods

### Strains

The collection of 33 *C. glabrata* strains used for the analyses in this study are listed in table S1.

### DNA extraction

*C. glabrata* cultures were grown overnight in an orbital shaker (200 rpm, 30 ºC) in 2 ml YPD (Yeast Peptone Dextrose) medium (0.5% yeast extract, 1% peptone, 1% glucose) supplemented with 1% penicillin-streptomycin solution (Sigma). Subsequently, cells were centrifuged (3000 rpm, 5 min) and washed twice with 1x sterile PBS. The pellet was resuspended in 500 μl lysis buffer (1 w/V% SDS, 50mM EDTA, 100 mM TRIS pH=8), afterwards 500 μl of glass beads were added to the cells which then were disrupted by using a vortex for 3 min. 275 μl 7M ammonium-acetate were added (65 ºC, 5 min) and the samples were cooled on ice for 5 min. Then 500 μl of chloroform-isoamylalcohol (24:1) were added to the mixture, which was then centrifuged for 10 min at 13000 rpm. The upper phase of the solution was transferred to a new microcentrifuge tube, and the previous step was repeated. 500 μl isopropanol was mixed with the upper phase of the solution in a new microcentrifuge tube, and the mixture was held in a refrigerator at −20 ºC for 5 min. The solution was centrifuged at 13000 rpm for 10 min. The supernatant was discarded, and the pellet was washed twice with 500 μl 70 % ethanol. After the second washing step the pellet was dried and resuspended in 100 μl bi-distilled water containing RNase (Sigma).

### Sequencing

The genome sequences for all the strains were obtained at the Ultra-sequencing core facility of the CRG, using Illumina HiSeq2000 sequencing machines. Paired-end libraries were prepared. For this, DNA was fragmented by nebulization or in Covaris to a size of ~600 bp. After shearing, the ends of the DNA fragments were blunted with T4 DNA polymerase and Klenow fragment (New England Biolabs). DNA was purified with a QIAquick PCR purification kit (Qiagen). 3’-adenylation was performed by incubation with dATP and 3’-5’-exo-Klenow fragment (New England Biolabs). DNA was purified using MinElute spin columns (Qiagen) and double-stranded Illumina paired-end adapters were ligated to the DNA using rapid T4 DNA ligase (New England Biolabs). After another purification step, adapter-ligated fragments were enriched, and adapters were extended by selective amplification in an 18-cycle PCR reaction using Phusion DNA polymerase (Finnzymes). Libraries were quantified and loaded into Illumina flow-cells at concentrations of 7–20 pM. Cluster generation was performed in an Illumina cluster station. Sequence runs of 2x100 cycles were performed on the sequencing instrument. Base calling was performed using Illumina pipeline software. In multiplexed libraries, we used 4 bp internal indexes (5’ indexed sequences). Deconvolution was performed using the CASAVA software (Illumina). All sequence data has been deposited in SRA and will be available upon publication.

### Genome assembly

Reads were pre-processed previous to assembly to trim at the first undetermined base or at the first base having PHRED quality below 10. The pairs with reads shorter than 31 bases after trimming were not included from the assembly process. SOAPdenovo2 (Luo et al. 2012) with default parameters was used to assemble paired-ends reads into chromosomes.

### SNP calling

Reads were aligned onto the reference assembly of the CBS138 strain (Dujon et al. 2004b) using BWA, with the BWA-MEM algorithm with 16 as number of threads (Li and Durbin 2010). As no raw reads are publicly available for the recently-sequenced pyruvate-producing strain *C. glabrata* CCTCC M202019 (Xu et al. 2016), the Wgsim software v0.3.1-r13 (Heng, Li) was used to simulate reads from the assembled genome sequence with by defaults parameters, with the exception of rate of mutations and fraction of indels that was 0 and number of read pairs that was set to 100000000.

We identified SNPs using GATK v3.3 (McKenna et al. 2010; DePristo et al. 2011; Van der Auwera et al. 2013) with an haploid model, filtering out clusters of 5 variants within 20 bases and lowquality variants, and using thresholds for mapping quality and read depth (>40 and >30 respectively). To confirm ploidy levels and assess heterozygosity in duplicated chromosomes we repeated the SNP calling analysis enforcing a diploid model. Thereafter variants were divided into homozygous and heterozygous categories.

### Structural variants

To detect structural variants we used deviations from the expected depth of coverage. Calling deletions or duplications at genomic regions with variable coverage is a widely accepted methodology (Boeva et al. 2011). For every C. *glabrata* strain we calculated the number of genes deleted and duplicated using depth of coverage analysis from Samtools (Li et al. 2009; Li 2011). After mapping the reads of each strain to the reference genome a gene was considered missing in the strain if less than 90% of the length of a given gene was covered by reads. For duplications and large scale structural variants, we normalized the number of reads per gene and a duplication was called if the median coverage of that gene was 1.8 times or higher than the median coverage of the chromosome. All these structural variants were manually curated and one deletion comprising three genes was validated experimentally (see below).

### Experimental validation of CNVs

The deletion of a region comprising three genes (corresponding to deletion numbers 33, 34 and 35 (Supplementary Table S2) was experimentally confirmed by means of PCR and Sanger sequencing in the following strains: EF1620Sto, F11, F15, EG01004Sto, F03013, F15021 and CBS138. Because the investigated fragment was 8,134 base pairs long, four different sets of primers were designed to be able to capture the whole fragment (Supplementary Figure S4, a). First PCRs were performed with CBS138 (control strain) and one of the investigated strains (EF1620Sto) and primers FWD1: REV1, FWD2: REV2, FWD3: REV3 to validate the absence of the deletion in the control strain and the feasibility of the primers and PCR reactions (Supplementary Figure S4, b). In this case amplicons from the three primer pairs are expected only in the control strain. Then the absence of the fragment was tested with the results of the PCR with FWD1: REV3 (Supplementary Figure S4, c). With this primer, the absence of a band in the CBS138 control strain indicates that the deletion is not present, and the amplification of a fragment of approximately 2 kbp long in the other strains confirms the existence of the deletion. DNA extraction was performed with the MasterPure Yeast DNA Purification Kit from EPICENTRE according to the manufacturer's protocol. PCRs were carried out by using *Pfu* DNA polymerase from PROMEGA. The reaction mixture included primer concentration of 0.4 µM, 5 µl of *Pfu* polymerase 10X buffer with MgSO4, 200 µM of dNTPs each, 1.2 U of *Pfu* DNA polymerase, 100 ng of DNA and water up to a final volume of 50 µl. Standard PCR protocol was used for primers: FWD1: REV1, FWD3: REV3 and FWD1: REV3. Here, initial denaturation was performed at 95°C for 2min, followed by 30 cycles of 30 sec at 92°C, 30 sec at either 60.3ºC (FWD1: REV1), 59.3ºC (FWD3: REV3) or 60.3ºC (FWD1: REV3); 190 sec (FWD1: REV1) or 120 sec (FWD3: REV3) or 220 sec (FWD1: REV3) at 72°C. It was finished with final extension for 5 min at 72°C and cooled to 4°C. The touchdown PCR was performed for FWD2: REV2. Cycling condition began with 2 min at 95°C, followed by 15 cycles of 30 sec at 95°C, 15 sec at the annealing temperature of 61.3°C (decreasing 0.5°C each cycle) and 4 min 20 sec at 72°C. Then, other 20 cycles of 30 sec at 95°C, 15 sec at the annealing temperature of 54.3°C and 4 min 20 sec at 72°C were set up, with a final extension step at 72°C for 5 min. All PCR products were visualized by 1% agarose gel electrophoresis (Supplementary Figure S4, b, c). and were then purified using the QIAquick PCR Purification Kit (QIAGEN) for subsequent Sanger sequencing. Sequences of the primers used (5'->3') were: FWD1: TTGGTCTGTTCCTGAGCCGG; FWD2: ACGAACTGGATAGCACCTCC, FWD3: ATACTGTGACCTTCCCTGTT; REV1: CTCAGCATTGGCAGTAGTGG; REV2: CTTCGCTCCGTGGGTAAACA and REV3:. CTTCAGATTGGCAGTGTCGG.

### PCR amplification of mating-type regions and sequencing

In order to validate mating-type switching in *C. glabrata*, we performed Sanger sequencing of the three different loci *MTL1*, *MTL2* and *MTL3* encoding for the two **a** and alpha mating types in 14 different strains: E1114, M6, CST110, EG01004Sto, F15021, F03013, BG2, P35_2, P35_3, M12, EI1815Blo, F11, EF1237Blo1 and the reference CBS138. DNA extraction and PCRs were performed as indicated in the previous section. Primers used (5'->3)' and expected amplicon sizes (bp) are as follow: MTL1_Forward: CGGTCTGATGGTGCAATTGT, MTL1_Reverse: TTGAGTCAAGTGTCGAGGCT (1760 bp); MTL2_Forward: GCTCTTCACTCAACGTACTCC, MTL2 _ Reverse: TTTACAAACCCACACCGAGG (1305 bp); MTL3 _ Forward: GTGAGCACTTTGGACCTTCA, MTL3_reverse: ACCATAGTCAGACCACCGAC (1908 bp). Briefly, each reaction included primer concentration of 0.4 µM, 5 µl of *Pfu* polymerase 10X buffer with MgSO_4_, 200 µM of dNTPs each, 1.2 U of *Pfu* DNA polymerase, 100 ng of DNA and water up to a final volume of 50 µl. Cycling condition began with a warm-up step of 2 min at 95ºC, followed by 30 cycles of 30 sec at 95ºC, 30 sec at the corresponding annealing temperature (55.5ºC, 58.4ºC and 58.3ºC for *MTL1*, *MTL2* and *MTL3*, respectively) and an elongation step at 72ºC for 3 min 50 sec, 2 min 20 sec, and 3 min 20 sec for *MTL1*, *MTL2* and *MTL3*, respectively, with a final elongation step of 72ºC for 5 min. PCR products were confirmed by 1.5% agarose gel electrophoresis, were then purified using QIAquick PCR purification kit according to manufacturer’s instructions (Qiagen) and finally sequenced with Sanger using the same set of primers.

### Recombination events

First, we select deletions that appeared in more than one clade. Second, for each selected deletion we extracted genomic regions located 1Kb up- and downstream of the affected gene. Then, we used RDP4 v4.15 (Martin and Rybicki 2000; Martin et al.) to identify footprints of homologous recombination, and to produce recombination rate plots (McVean et al. 2004). Finally, we used SplitTree v4(Huson and Bryant 2006) to construct phylogenetic networks and to detect gene flow.

### Chromosomal rearrangements

We identified the presence of chromosomal arrangements using two steps. First, we obtained *denovo* assemblies of all genome from the 32 *C. glabrata* strains. Second, we reordered them using CBS138 as a reference applying Mauve Contig Mover from Mauve (Darling et al. 2004).

### Phylogenetic analysis

We reconstructed a species tree including the 32 sequenced *Candida glabrata* strains, the reference strain CBS138 and *Candida bracarensis* (CBS10154) (Gabaldón et al. 2013) as the outgroup. By using the previously annotated SNPs for the 32 strains, we reconstructed the sequence of each strain by replacing the reference nucleotide for a given SNP. Then, these 34 genomes were aligned using Mugsy v1.2.3 (Angiuoli and Salzberg 2011). The resulting alignment was trimmed using TrimAl v.1.4 (Capella-Gutiérrez et al. 2009) to delete positions with more than 50% gaps. Finally a phylogenetic tree was reconstructed from the trimmed alignment using RAxML v7.3.5, model Protgammalg(Stamatakis et al. 2005).

### Population Genomics

We used the software STRUCTURE v2.3.4 to study the genetic structure of the population (Hubisz et al. 2009). In addition we used popGenome to estimate FST between different clades (Pfeifer et al. 2014). We recorded the number of SNPs in *C. glabrata* population using the 33 strains. We obtained the number of SNP also using 32 different strains from *C. albicans,* as indicated above for *C.glabrata,* and we obtained SNP data from *Saccharomyces cerevisiae* using dbSNP(Sherry et al. 2001) as of October 2015. We calculated the ratio of non-synonymous and synonymous nucleotide diversity (πN/πS) assuming that ¾ of all of the sites are non-synonymous, and ¼ synonymous.

### Phenotypic analyses (Growth curves)

Each strain was recovered from our glycerol stock collection and grown for 2 days at 37 ºC on a YPD agar plate. Firstly, single colonies were cultivated in 15 ml YPD medium in an orbital shaker (37ºC, 200 rpm, overnight). Secondly, each sample was diluted to an optical density (OD) at 600 nm of 0.2 in 3 ml of YPD medium and grown for 3 h more in the same conditions (37ºC, 200 rpm). Then, dilutions were made again to have an OD at 600 nm of 0.5 in 1 ml of YPD medium in order to start all the experiments with approximately the same amount of cells. The samples were centrifuged for 2 min at 3000 g, washed with 1 ml of sterile water, and centrifuged again for 2 min at 3000 g for a final resuspension of the pellet in 1 ml of sterile water. Finally, 5 µl of each sample was inoculated in 95 µl of the corresponding medium in a 96-well plate. All experiments were run in triplicate.

A total of six different growth conditions were tested: the oxidative stress was assessed by the growth of the cultures on YPD medium supplemented with 10 mM H2O2, reductive stress with 2.5 mM DTT and osmotic stress with 1 M NaCl. We also measured the impact of elevated temperature (41.5ºC), pH=2 and pH=9 along with the control growth on YPD itself. Cultures were grown in 96-well plates at 37ºC or 41.5ºC, shaking, for 24 or 72 h depending on the growth rate in each condition, and monitored to determine the optical density at 600 nm every ten min by a TECAN Infinite^®^ M200microplate reader.

### Phenotypic analyses (biofilm formation assay)

The capacity to form biofilms was assayed as described before (Gómez-Molero et al. 2015a). Briefly, studied isolates and controls (CBS138, moderate biofilm formation capacity; PEU-382 and PEU-427, high biofilm formation capacity were cultured overnight in YPD medium at 37°C. The optical density was determined at 600 nm (Ultrospec 1000) and adjusted to a value of 2 using sterile NaCl_physiol_. 50 μl aliquots of the cell suspensions were placed into 96-well polystyrol microtiter plates (Greiner Bio-one) and incubated for 24 h at 37°C. The medium was removed and the attached biofilms washed once with 200 μl distilled water. Cells were stained for 30 min in 100 μl of 0.1% (w/v) crystal violet (CV) solution. Excess CV was removed and the biofilm carefully washed once with 200 μl distilled water. To release CV from the cells, 200 μl 1% (w/v) SDS in 50% (v/v) ethanol were added and the cellular material resuspended by pipetting. CV absorbance was quantified at 490 nm using a microtiter plate reader (MRX TC Revelation). The data shown is the average of three independent biological experiments, each including four technical repeats.

### Antifungal drug susceptibility testing

Prior to analysis, the isolates were cultured overnight on Sabouraud (Oxoid) agar plates. Antifungal drug susceptibilities towards Fluconazole, Isavuconazole, Posaconazole, Voriconazole, Micafungin, Caspofungin, 5-Fluorcytosine, and Amphotericin B were determined according to EUCAST EDef 7.1 method (Arendrup et al. 2012). The MIC values of each isolates were calculated according to EUCAST guidelines (http://www.eucast.org/fileadmin/src/media/PDFs/EUCAST_files/AFST/Clinical_breakpoints/Antifungal_breakpoints_v_8.0_November_2015.xlsx, accessed Nov 16^th^ 2016)

### Pulsed-field gel electrophoresis

Intact chromosomes were separated using pulsed-field gel electrophoresis (PFGE) as described before (Bader et al. 2012). To better visualize differences between small chromosomes (size range CBS 138 ChrA-K) conditions were modified to using 1.2% agarose at 17°C and pulse times from40-100 sec. Large chromosomes (size range CBS138 ChrK-M) were resolved with pulse times form 60-140 sec.

### Statistical analyses and plots

Fisher tests were computed using R and plots were obtained using the ggplot2 package for R (Wickham 2009). Multiple Correspondence Analysis (MCA) was performed using ade4 package for R to establish the main relationships between all sequenced strains and the reference (Tenenhaus and Young 1985). MCA is a technique similar to principal component analysis (PCA) but specific for nominal categorical data, which is used to detect and represent underlying structures in data sets.

### Data Access

Sequence data produced for this project has been deposited at short read archive under the accession PRJNA361477

## Acknowledgements

TG group acknowledges support of the Spanish Ministry of Economy and Competitiveness grants, ‘Centro de Excelencia Severo Ochoa 2013-2017’ SEV-2012-0208, and BFU2015-67107 cofounded by European Regional Development Fund (ERDF); from the European Union and ERC Seventh Framework Programme (FP7/2007-2013) under grant agreements FP7-PEOPLE-2013-ITN-606786 “ImResFun” and ERC-2012-StG-310325; from the Catalan Research Agency (AGAUR) SGR857, and grant from the European Union’s Horizon 2020 research and innovation programme under the Marie Sklodowska-Curie grant agreement No H2020-MSCA-ITN-2014-642095. CF and TG's groups acknowledge support from the GDRI "iGenolevures" of the French CNRS, for travel and meeting funds. OB and EGM acknowledge funding from the European Union under grant agreement FP7-PEOPLE-2013-ITN-606786 “ImresFun”. All authors acknowledge the technical support of the UPF-CRG FACS facility and the CRG Genomics facility.

## Supplementary Figures and Tables

### Supplementary tables

### Supplementary Table S1

Information about the 33 *C. glabrata* isolates analyzed in the present study (including reference CBS138). Columns indicate, in this order: Strain name or ID; Synonym (if any); Mean sequencing coverage (if sequenced in this study); Body site of isolation; Country of isolation; Mating type; CC (clonal complex); RT (repeat type); Publication describing the source. “*” near strains name indicates commensal strain. NA indicates “not assigned”.

### Supplementary Table S2

Genes affected by CNVs. Columns indicate, in this order: Duplication or deletion number in this study (corresponds to Figure 1); Gene ID; Gene Name; Name of *S. cerevisiae* one-to-one ortholog (if any); description.

### Supplementary Table S3

Information about the 32 strain of *Candida albicans* re-analyzed in this study. Columns indicate, in this order: Strain name or ID; Host; Isolation site; Country of isolation; Mating type (if any); Experiment name in SRA; Number of SRA run.

### Supplementary Table S4

Synonymous vs non-synonymous variation per gene. First column indicate gene functional category (based on their function in *S. cerevisiae*). Following columns indicate gene ID, gene name (only for *S. cerevisiae*), and πN/πSfor *C. glabrata, C. albicans* and *S. cerevisiae* genes.

### Supplementary Table S5

Antifungal sensitivity. Range of sensitivity levels (triplicate experiment) to Amphotericin B (MIC90), 5-Fluorcytosine, Fluconazol, Voriconazol, Posaconazol, Isavuconazol, Micafungin, and Caspofungin (all MIC5 0 and, where available (www.eucast.org/breakpints) classification according to clinical breakpoints. S(Sensitive)/I(Intermediate)/R(Resistant). Red and boldfaced names and values, indicate isolates with MIC values measured deviating from the wild type distribution. There are no species-specific clinical breakpoints available for 5FC, Voriconazol, Posaconazol, Isavuconazol, or Caspofungin. Measured MIC range in most isolates (three technical repeats) encompasses both “susceptible” and “resistant” interpretations according EUCAST clinical breakpoint definition: S≤0.032, R>0.032 for Micafungin.

### Supplementary Table S6

Private mutations per strain (non-synonymous SNPs and CNVs). Columns in the tables of private non-synonymous SNPs indicate, in this order: strain ID; Chromosome and position affected by the SNP; Gene ID; Amino acid substitution; Gene name; description. First column of table based on CNV indicates strain analyzed. Following columns indicate: Gene affected, Description of gene affected.

### Supplementary Table S7

Levels of nucleotide variation per strain, mutations in MSH2. Columns indicate, in this order: Strainname; Clade; Non-synonymous variant; Saccharomycotina species presenting the same non-synonymous mutations. Five-letter codes corresponds to species mnemonic based on Uniprot taxonomy database (Pundir et al., UniProt Consortium 2015); Reduced sensitivity of the strain to any antifungal tested; genome-wide π_N_; genome-wide π_S_; genome-wide π_N_/π_S_.

## Supplementary Figures

### Supplementary Figure S1

Profile of SNPs densities obtained when comparing the genomes of strains in Clade I and Clade II using non-overlapping 10Kb windows along the entire genome. Bar on the top indicates order and relative length of C. glabrata chromosomes in CBS138. First profile indicates SNP density between the two strains from Clade I (M7 versus B1012M). Second and third profiles indicate SNP density between EB0911Sto and CST35 from Clade II versus Clade I (using B1012M as a reference for this clade). Forth profile indicates SNP density between the two strains of Clade II.

### Supplementary Figure S2

Heatmap showing average pairwise differences (SNP/Kb) between all strains analyzed in the present study. Strains are grouped by clades designated with different colors as in Figure 1.

### Supplementary Figure S3

3D scatterplot of the multiple correspondence analysis (MCA) with overlaid information regarding mating type (a), isolation location (b), body-site source (c), and clade distribution (d).

### Supplementary Figure S4

PCR results for the experimental confirmation of a deletion affecting three genes (corresponding to DEL nº 33, 34 and 35, in Figure 1, Supplementary Table S2). a) Scheme of the genomic context, the position of the primers and length of the amplicons in reference *C. glabrata* CBS138. b) PCR validation of the deletion in EF1620Sto and the absence of the deletion in the *C.glabrata* CBS138.c) PCR validation of the presence of the deletion in the tested strains.

### Supplementary Figure S5

Analysis of biallelic SNPs per strain when variant calling was performed using an haploid (top) or diploid (bottom) model. A similar fraction of biallelic SNPs as the one found in a control strain known to be haploid (BG2) suggests all strains are haploid.

### Supplementary Figure S6

Flow cytometry analyses of DNA content of *Candida glabrata* samples. Eight different *Candida glabrata* strains plus one *Candida* diploid sample (marked with an asterisk in the figure) were analyzed using the flow cytometry analyzer FACS LSRII. In the histogram, FITC-A values corresponding to DNA content *versus* cell counts are plotted using FlowJo software. For haploid samples, peaks of FITC-A around 17-28 K and 41-50 K account for cells in G1 and G2 phases, respectively, while the positive control diploid sample shows the FITC-A peak corresponding to cells in G1 and G2 phases at 50 K and 97 K, respectively. In the table, FITC-A G1 and G2 medians are shown per each analyzed strain, additionally with the ratio between them. The final column of the table shows the ratio between the G1 peak of each sample in comparison to the reference CBS138: all samples have a ratio around 1 (0.91-1.52) except for SZMC 8029, which is 2.70, confirming that it is the only diploid sample among all the ones analyzed.

### Supplementary Figure S7

Electrophoretic karyotyping (Pulse Field Gel Electrophoresis). Sizes of chromosomes are indicated for the reference strain CBS138 (rightmost lane).

### Supplementary Figure S8

Plots showing the aneuploidies detected in the present study. Plots indicate m ean coverage in bins containing 10 consecutive genes each. Data is shown for the four strains showing aneuploidies (F15021 EF0616Blo1, F1822, I1718), and B1012M as a control without aneuploidies. Three plots correspond to three chromosomes. From left to right: Chromosome B, as control without aneuploidies; Chromosome E (duplicated in EF0616Blo1, F1822, I1718); and Chromosome G with a partial aneuploidy in F15021.

### Supplementary Figure S9

Analysis of biallelic SNPs per chromosome when variant calling was used with the haploid (top) or diploid (bottom) model. The four strains with aneuploidies are shown, the aneuploid chromosome is indicated with an asterisk.

### Supplementary Figure S10

Spontaneous duplication of chromosome J in strain F2229. The distribution of the depth of coverage per gene in each of the chromosomes is plotted in first (left) and second (right) sequencing rounds. The left plot shows an increase in depth of coverage in Chromosome J. However this is only 1.5 higher than the coverage in the other chromosomes (a duplication would be expected to cause doubling of the coverage), indicating that approximately only 50% of the cells carry the aneuploidy. A second round of sequencing started from a fresh culture of the stored strain shows no chromosome with increased coverage, indicating that the previous aneuploidy appeared spontaneously during growth.

### Supplementary Figure S11

Recombination rate plot (left) and NeighborNet splits network (right) for genes showing signs of recombination. (Left) Each plot shows the average recombination rate (rho/bp) for the 33 strains computed for each gene and 1kb up- and downstream. Positions with rho/bp upper than 0.05 indicates a hotspot recombination site. Recombination was estimated using the LDhat method as implemented in the RDP4 software. (Right) NeighborNet splits network showing gene flow between strains. Clades are indicated as dots in different colors. Note that for some genes, strains of different clades are clustered together, suggesting a recombination event. NeighborNet networks were computed using Splits Tree software.

### Supplementary Figure S12

PCR amplification of mating-type regions. Amplicons obtained by means of PCR per each of the mating-type regions *MTL1*, *MTL2* and *MTL3* are shown in panel a, b and c, respectively. A schema with the genomic context and the primer position for each amplified region is shown in the top of each panel.

### Supplementary Figures S13

Phenotype analysis in *C. glabrata* testing growth rate using 7different conditions. First condition was YPD as a normal medium to growth. Following conditions were H2O2, NaCl, DTT, high temperature (41.5 Tº), basic pH (pH=9) and acid (pH= 2). Unless indicated otherwise all growth curves were carried out at 37ºC. The y axis shows the OD for each clade and the x axis shows time (in minutes). Supplementary Figure divided in seven parts, one for each clade starting with Clade I and finishing with Clade VII.

### Supplementary Figure S14

Biofilm formation test. Results represent averages of three independent replicas of four technical repeats each. Positive controls are well characterized clinical isolates from urine and respiratory material, respectively, with known high adherence phenotypes(Gómez-Molero et al. 2015b).

